# Functional diversities within neurons and astrocytes in the adult rat auditory cortex revealed by single-nucleus RNA sequencing

**DOI:** 10.1101/2024.04.16.589831

**Authors:** Aysegul Gungor Aydin, Alexander Lemenze, Kasia M Bieszczad

**Author notes:** **Corresponding author** Correspondence to Dr. Kasia M. Bieszczad, 327 Psychology Building, Rutgers University—New Brunswick, 152 Frelinghuysen Road, Piscataway, New Jersey 08854 USA.

## Abstract

The mammalian cerebral cortex is composed of a rich diversity of cell types. Cortical cells are organized into networks that rely on their functional diversity to ultimately carry out a variety of sophisticated cognitive functions. To investigate the breadth of transcriptional diverse cell types in the sensory cortex, we have used single-nucleus RNA sequencing (snRNA-seq) in the auditory cortex of the adult rat. A variety of unique excitatory and inhibitory neuron types were identified. In addition, we report for the first time a diversity of astrocytes in the auditory cortex that may represent functionally unique subtypes. Together, these results pave the way for building models of how neurons in the sensory cortex work in concert with astrocytes at synapses to fulfill high-cognitive functions like learning and memory.

## Introduction

The mammalian cerebral cortex carries out a variety of sophisticated cognitive functions through organized networks of diverse cell types^1–6^. Specific cell types differ in connectivity and physiological properties established in part by their transcriptional profiles. For example, subtypes of sensory gamma-aminobutyric acid-ergic (GABAergic) interneurons are identifiable by gene products that produce unique molecular markers (e.g., *Gad1*) and have specific contributions within cortical circuits for modulating sensory processing and information coding^7–14,5,15^. Sensory cortical areas are characteristically dynamic in support of task- and experience-dependent processes^16–20^. This characteristic is afforded in part by the flexibility of changes to cellular function within an assortment of potential cellular identities that can be activated by behavioral demands. The potential for the dynamics within cortical circuit function may be key to ultimately support physiological plasticity for adaptive sensory information processing across a lifetime^16,21–24^. The auditory cortex (AC) is a well-known site for task- and experience-dependent neurophysiological plasticity and one of the main models for understanding substrates of auditory learning and memory^23,25^, but its basic transcriptomic landscape is under-described. Characterizing gene expression profiles of the diverse range of auditory cortical cell populations can inform on their potential functions, interactions, and our understanding of how the AC supports auditory processing, learning, memory, and sound-driven behavior. Here, we provide a transcriptomic foundation for understanding the cellular and functional physiological substrates of higher-level cognitive processing in rats, a species well-known for its natural curiosity and intelligence, bolstered by an enormous behavioral literature^26–29^ and its particular use to investigate memory, cognition, and complex behavior^23,25,30–33^.

## Results

### Identification of transcriptomic profile of cell types in adult rat AC

To generate a multicellular map of gene expression at single-nucleus resolution in the AC, we performed single-nucleus RNA sequencing (snRNA-seq) and analyzed 255,000 total nuclei from 2 adult male rats (Sprague-Dawley). AC tissue was dissected from flash-frozen brains, bilateral hemispheres were combined, nuclei were isolated (see *Methods*) and then captured, and their mRNA was barcoded using the split-pool barcoding approach^34^. After data preprocessing and quality control, we acquired 11,952 high-quality single-nucleus transcriptomes with a median of 2607 unique molecular identifiers (UMIs) and 1432 genes per nucleus.

First, to identify cell subpopulations in rat AC, we used unsupervised dimensionality reduction approaches to classify individual cells into cell types according to their gene expression patterns. We plotted the clusters using Uniform Manifold Approximation and Projection (UMAP) to discover molecularly distinct classes of cells and identified 15 discrete cell clusters, excluding one small cluster of nuclei derived from the surrounding hippocampus, which was removed from subsequent analyses (see *Methods*) (**Fig. 1A** and **Supplemental Fig. 1**). Next, we used canonical marker genes for excitatory and inhibitory neurons (e.g., *Rorb* and *Gad1*), as well as for astrocytes (e.g., *S100b*), oligodendrocytes (e.g., *Mog*), oligodendrocyte precursor cells (e.g., *Pdgfra*), and microglia (e.g., *Cx3cr1*) to annotate each cell clusters.

**Figure 1.**
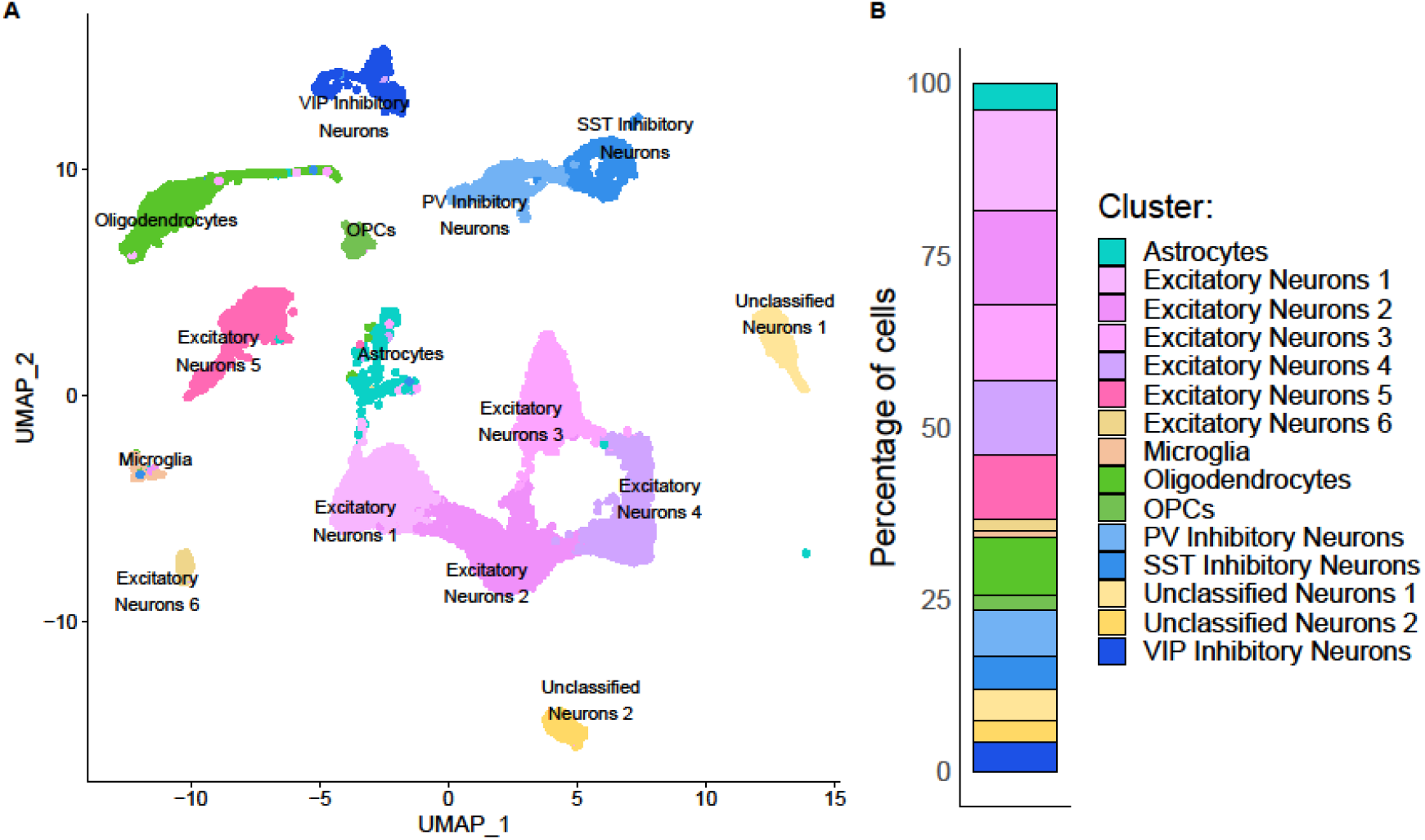
Adult rat AC single-nuclei transcriptomic data. **(A)** Uniform Manifold Approximation and Projection (UMAP) visualization of clusters with all major identifiable cell types labeled. Dots represent gene expression profiles for individual nuclei and are colored by cell type. OPCs, oligodendrocyte progenitor cells. **(B)** Quantification of the percentage of different cell types revealed by single-nucleus RNA sequencing (snRNA-seq) datasets in rat AC.

Based on the expression of canonical markers, of the 15 clusters, 13 could readily be labeled using standard markers. These 13 clusters classified six major cell types with varied proportions of inhibitory neurons (1863 nuclei, 15.9% of all nuclei), excitatory neurons (7199 nuclei, 61.3% of all nuclei), oligodendrocytes (990 nuclei, 8.4% of all nuclei), oligodendrocyte precursor cells (OPCs) (246 nuclei, 2.1% of all nuclei), astrocytes (446 nuclei, 3.8% of all nuclei), and microglia (103 nuclei, 0.87% of all nuclei) (**Fig. 1**). Two clusters, unclassified neuron 1 and 2, could not unequivocally be labeled with standard markers (“Unclassified Neurons 1” and “Unclassified Neurons 2” in **Fig. 1A&B**). These neurons are mostly GABAergic but also heterogeneous. Indeed, we also found putative interneuron marker genes *Sema3c*^35,36^ and *Slit2*^36^ in the first, and *Kcnc2*^37^, *Sox5*^38^ were expressed in the second unclassified cluster. On the other hand, we also found expression of layer-specific excitatory neuronal marker genes such as *Tox*^36,37^ (L5) in unclassified neuron 1 as well as *Cux2*^37,39^ (L2-4) and *Nr4a2*^37^ (L6B) in unclassified neuron 2. This subset of neurons in the deeper cortical layer 6B introduces intriguing potential inhibitory influences on the subcortical outputs of AC and requires further investigation of their putative functional relevance.

### Cellular and molecular profiling of inhibitory and excitatory neurons

Next, to uncover molecular diversity within inhibitory and excitatory cell types, cells classified within one of the main cell types were isolated and re-clustered to delineate cellular subpopulations (see *Methods*). Clustering analysis of inhibitory neurons (a total of 1863 nuclei) revealed three transcriptomic cell types belonging to three previously described^40–42^ major subtypes of GABAergic inhibitory neurons in AC: Vip+ (vasoactive intestinal peptide), Pv+ (parvalbumin), and Sst+ (somatostatin) (**Fig. 2A**). In accord with previous reports, these subtypes are defined by prominent or combinatorial expression of marker genes *Sox5* for Pv^38^; *Grin3a* for Sst^35^; *Vip* and *Adarb2*^6,43,44^ for Vip inhibitory neurons, along with pan and putative interneuron markers *Gad1*^5,43,45^ and *Erbb4*^37,38,43,46^ (**Fig. 2C, left**). Each inhibitory subtype appeared in varying proportion in this dataset; Pv inhibitory neurons (795 nuclei, 42.7% of inhibitory neurons), Sst inhibitory neurons (561 nuclei, 30.1% of inhibitory neurons), and VIP inhibitory neurons (507 nuclei, 27.2% of inhibitory neurons) (**Fig. 2C**, right).

**Figure 2.**
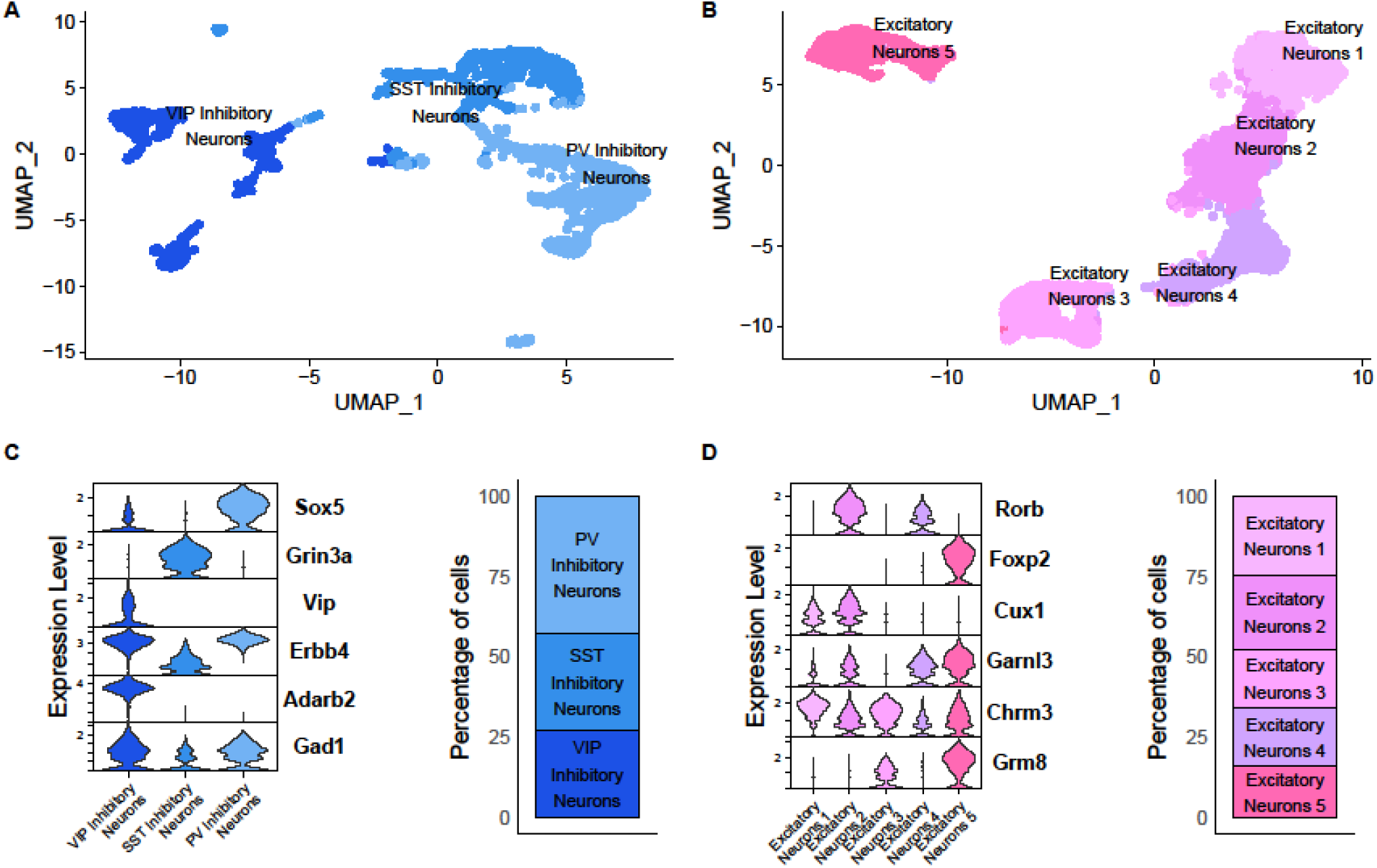
Inhibitory and excitatory neuron subtypes and selected marker genes. **(A)** Annotated UMAPs of inhibitory (n = 1863 nuclei) and **(B)** excitatory neuron subtypes (n = 6997 nuclei). Colors indicate clusters identified by graph-based clustering. The violin plot represents the distribution of subtype-specific marker genes (left) and the proportion of each subtype (right) for **(C)** inhibitory and **(D)** excitatory neurons. Rows are genes, and columns are cell types. Values within each row are SCT-normalized.

For excitatory cells (a total of 6997 nuclei), we identified five transcriptomic subtypes of excitatory neurons (**Fig. 2B**), each showing enrichment of specific genes (**Fig. 2D, left**). These subtypes appeared to be broadly segregated by layer by the expression of known cortical lamina molecular markers. Examples of layer-specific genes include *Rorb*, a marker of L4; *Foxp2*, a marker of L6; and *Cux1*, a marker of L2-4, consistent with the previous publications^6,39,47,48,48,49^. *Cux1* and *Rorb* were enriched in Excitatory neurons 1 & 2, and 2 & 4, respectively. However, *Foxp2* was restricted to Excitatory neurons 5 (**Fig. 2D, left**). Excitatory neuron subtypes constituted relatively similar proportions of the total cell population analyzed, ranging from 16% (Excitatory Neuron 5) to 24.7% (Excitatory Neuron 1).

In addition to the five prominent excitatory neuron populations described in **Figure 2**, we also identified one small excitatory neuron cluster, Excitatory Neuron 6, which was most probably derived from cells of the subiculum (**Fig. 1A**). While these cells expressed deep-cortical layer excitatory gene markers such as *FoxP2*^50^ (L6a), *Tox*^36,37^ (L5), and *Tle4*^37^ (L6), cells in this cluster co-expressed genes are known to mark the subiculum such as *Olfm3*^36,51^, *SSbp2*^51,52^, *and Neto2*^36,51^. To be conservative in our analysis, we excluded the Excitatory Neurons 6 cluster from subsequent re-clustering analyses, as it could not be confidently ascribed to AC. This neuron cluster is likely to represent technical limitations during tissue punching as the close proximity between the AC and surrounding subcortical regions may increase the likelihood of capturing these cell types together.

### Identification of transcriptionally distinct astrocyte subtypes

We next sought to explore the heterogeneity of characteristic astrocytes in the AC. To do so, all other cell types were removed to re-cluster a total of 446 astrocyte nuclei (see *Methods*). This led to the identification of four distinct Astrocyte SubTypes (AST1–4), each expressing a unique transcriptomic profile (**Fig. 3A**). Different ASTs showed enrichment of specific genes: *Sox6 and Nckap5* (AST1); *Kcnip4 and Nrg3* (AST2); *Fn1 and Tek* (AST3); and *Syp and Celsr2* (AST4) (**Fig 3B, right**). ASTs constituted very different proportions of the total cell population, ranging from 4.3% (AST4) to 38.8% (AST1) (**Fig. 3B, right**).

**Figure 3.**
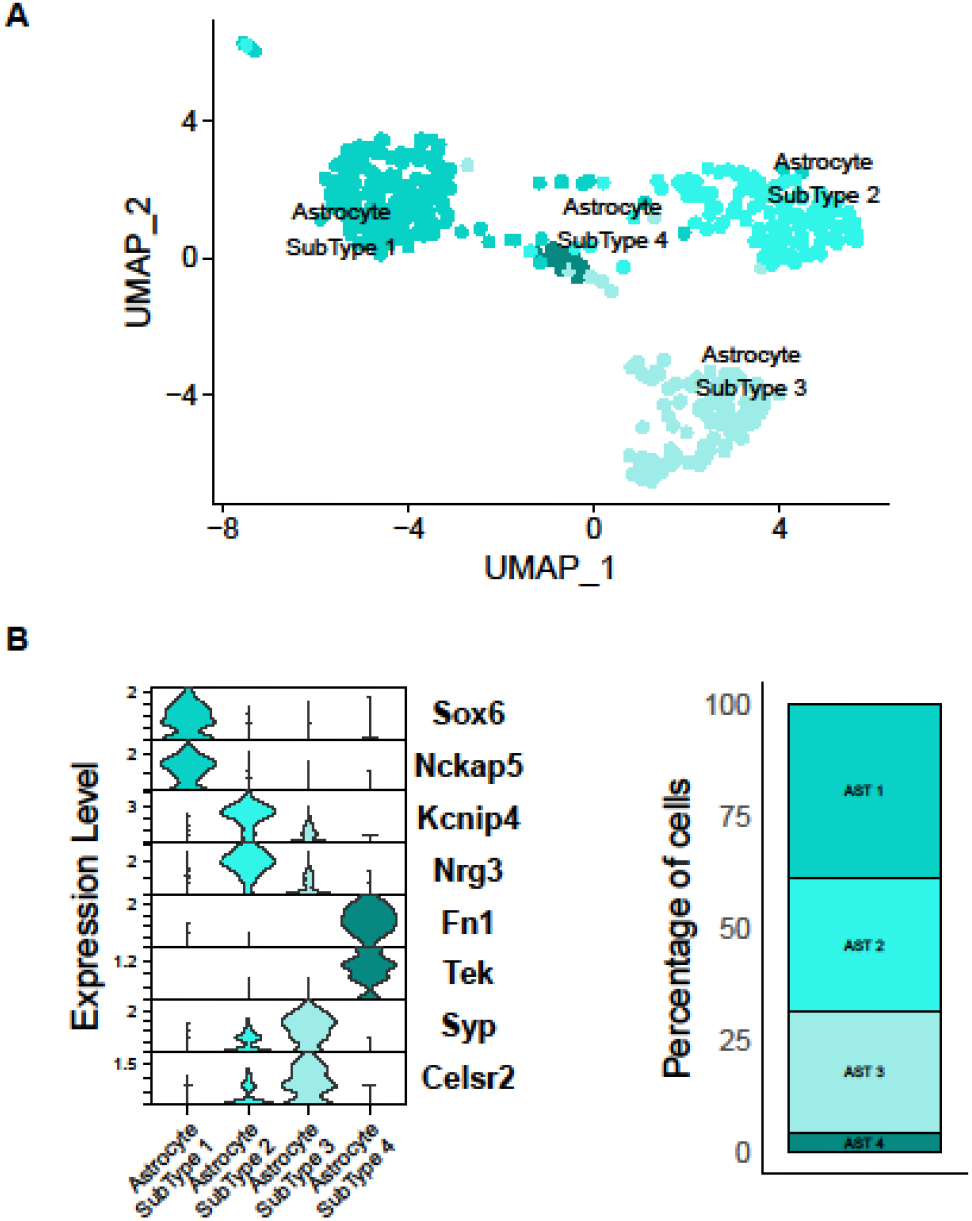
Identification of astrocyte subtypes in adult rat AC. **(A)** UMAP visualization of adult rat AC astrocytes (n = 446 nuclei). **(B)** The violin plot represents the distribution of subtype-specific marker genes (left) and the proportion of each subtype (right) for astrocyte sub-populations. Rows are genes, and columns are cell types. Values within each row are SCT normalized. values.

### Functionally diverse astrocyte subtypes

Finally, to begin to understand the potential functional identity of each AST, we examined differentially expressed genes (DEGs) using Gene Ontology (GO) enrichment analysis. We identified uniquely or commonly expressed genes of the identified subtypes and found three genes commonly expressed in each AST (**Fig. 4A**). In contrast, hundreds of other genes were differentially expressed between AST1–4. Interestingly, AST2 and AST3 have the largest number of overlapping DEGS (137) suggesting their functional similarity. For example, both subtypes include genes *Kcnd2* (ion transport), *Chrm3* (neurotransmission), *Ndrg4* (synapse function/plasticity), and *Erc2* (synaptic assembly) (**Fig. 4A**). Indeed, we found that AST2 and AST3 were highly enriched for genes associated with synapse organization and function, with an over-representation of biological processes such as synapse assembly, regulation of synapse structure and activity, and synaptic transmission (**Fig. 4B**). GO analyses revealed that the biological enrichment of DEGs between two other subtypes, AST1 and AST4, were primarily related to the angiogenesis and vasculogenesis (AST4), or myelination and oligodendrocyte function (AST1). Importantly, this pair appeared fundamentally different from the first pair of astrocyte subtypes (AST2 and AST3). The latter may have particular functional involvement in synaptic regulation, possibly through a tripartite synaptic relationship between pre- and post-synaptic neurons^53,54^. These two astrocyte subtypes are intriguing for their potential influences on the dynamics of neurophysiological plasticity in AC and are now primed for future investigation of their putative functional relevance in task- and experience-dependent processes.

**Figure 4.**
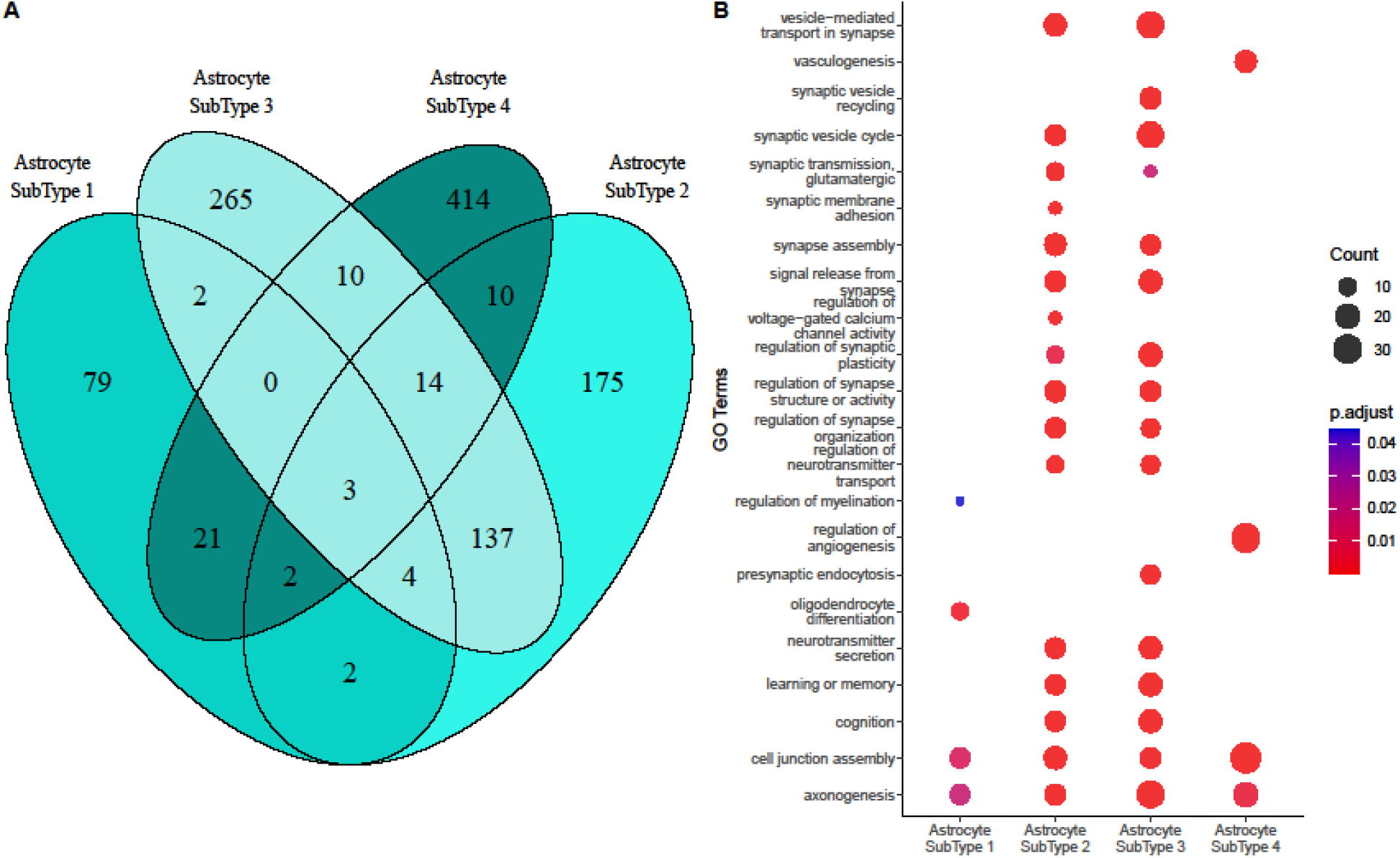
Comparison of astrocyte subtypes (ASTs)-specific transcriptomic profile in the AC. **(A)** Venn diagrams of the number of unique and shared Gene Ontology (GO) terms in four ASTs. **(B)** The bubble plot shows selected pathway enrichment results from DEGs among ASTs. Each row indicates a Gene Ontology (GO) biological process, and each column indicates the AST from which the DEGs are obtained. The bubble size represents the number of genes comprising each GO biological process. Color represents statistical significance, indicated by the legend on the right (adjusted p-value).

## Discussion

Knowledge of the specific interneuron populations that subserve distinct modes of inhibitory control have recently provided key insights for modulating information processing under specific behavioral conditions in the sensory cortex^55–62^. For example, activating layer 1 (L1) ionotropic 5-HT3A receptor (5-HT3AR) positive inhibitory interneurons in rodent primary auditory cortex (A1) resulted in distinct inhibition of parvalbumin-expressing (PV) and pyramidal cells in L4 to orchestrate critical-period plasticity in A1^55^.

Inhibition shapes cortical circuit activity^63,64^ and sensory processing^7,11,62,65^ through highly specific and precise spatiotemporal inhibitory control of the activity of glutamatergic excitatory neurons and local networks^10,66,67^ during both development^55,68,69^ and adulthood^70,64^.

Our data showed good overall coverage of different inhibitory and layer-specific excitatory neurons in adult AC. For example, we found fast-spiking interneurons called chandelier cells (PV interneuron marker ErbB4^38,71^), which has been implicated in synaptic plasticity by sensory experience in the visual cortex^72,73^. Also, we captured *Garnl3* expressing glutamatergic corticothalamic projection neurons^38^. In addition to these well-studied neurons, the data here provide new information about additional populations that we highlight by the top 5 DEGs within clusters: *Tafa1, Cntn6, L3mbtl4, Sema3c*, and *Vat1l* for unclassified neuron 1; *Zfp804b, Ttn, Shisa6, Ntng2*, and Lypd6 for Unclassified Neuron 2. These cells are now subject for further studies to reveal their potential functional importance.

A growing body of literature implicates the significance of inhibitory tone on excitatory cortical networks and opens the door to considering how other cell types could modulate cortical circuits. There is a mounting recognition that astrocytes play a significant role in synaptic modulation of circuits and behavior in the adult brain^74–76^. It is important to consider that glial cells outnumber principal cells by nearly 2:1^77^. We successfully identified multiple subtypes of astrocytes (AST1—4) at the transcriptomic level from a relatively low number of cells compared to principal cells. Consistent with our results, recent cortical studies also report astrocyte subpopulations, e.g., in somatosensory areas^78,79^. The findings of unique transcriptomic profiles of astrocytes highlight several intriguing possibilities, including functional specializations within known astrocyte function^75,80,81^. Furthermore, the number of unique genes between ASTs outnumbered the shared ones, which indicates potentially uniquely specialized astrocyte cell types^76,82,83^. For example, AST4 possesses a unique transcriptome with a small overlap with other ASTs, and GO terms suggest that this subtype could be linked to functions in the neuro-vascular unit^80^. However, the large overlap between AST2 and AST3 in DEGs show similar enrichment patterns for specific terms, suggesting their similar functional roles. The most common GO terms shared between these subtypes were involved in synapse assembly, regulation of synapse organization, signal release from the synapse, regulation of synapse structure or activity, vesicle−mediated transport in the synapse, synaptic neurotransmission, and a few others. Overall, this report highlights how transcriptomics can reveal a richer diversity of adult AC cell types and encourage consideration of a more intricate picture of canonical cortical circuitry. Together, these sensory cortical cells serve perception, cognition, and behavior, and their variety may be particularly useful in dynamic sensory environments and in a variety of task- and experience-dependent challenges. Considering the involvement of astrocytes in information processing through tripartite synapses^53,74,84^, it is crucial to reveal potential causal relationships between transcriptomic signatures and for a better understanding of cortical structure and function.

Thus, our findings suggest new directions for further investigation. At the forefront is the question of the relationship between transcriptomic signature and functional roles. For example, what specific roles and functions do astrocyte subclusters that have synaptic-related GO terms, serve within auditory neural circuits, and how does this heterogeneity contribute to the specificity of astrocyte control of neural circuits and behavior? Do the transcriptomic profiles of these astrocytes change in an activity-dependent manner? More excitingly, can we target these astrocytes to promote synaptic and neuronal functions or neuroplasticity for sound-driven behavior? Although these questions await further studies, our short report provides foundational insights into the transcriptional landscape of the adult AC and deciphers the underlying cellular and molecular underpinning of cortical circuits that regulate the flow of information and produce auditory cognition.

In summary, our results revealed a remarkable heterogeneity among astrocytes in AC. This diversity is characterized not only by transcriptomic differences but also by distinct functional roles encoded in their gene signatures, shedding light on the multifaceted roles of astrocytes in auditory circuits for higher-level cognitive processing. Our findings may be a valuable resource and a highly promising entry point for a deeper understanding of astrocytic roles in auditory processing, learning, and memory.

## Methods

### Subjects

Adult male Sprague-Dawley rats (n=2, 275-300g on arrival) were purchased from Charles River Laboratories (Wilmington, MA) and were individually housed in cages with various enrichment materials added, including shredded paper nesting material, gnawing block, and plastic shelter in a colony room with a 12 h light/dark cycle. Rats were water-restricted, with daily supplements provided to maintain them at ∼85% free-drinking weight and were briefly handled daily prior to tissue collection (1-2min/day) to acclimate them to the experimenters. All procedures were approved and conducted in accordance with guidelines by the Institutional Animal Care and Use Committee at Rutgers.

### Tissue collection from adult AC

Rats were deeply sedated with isoflurane and then euthanized by decapitation. Brains were rapidly removed and flash-frozen in pre-chilled 2-methyl-butane for 60 sec. Frozen brains were stored at −80 C until sectioning. Flash-frozen brains were embedded in an optimal cutting temperature medium (OCT), sectioned into 250-um-thick horizontal sections at −20 °C with the Leica (CM 3050S) Cryostat, and placed on a glass slide. ∼ 12 micropunches/sample (from both hemispheres) were obtained for single nuclei isolation from the 250-μm sections containing the AC (ranging from −5.82 to −3.86 anterior-posterior (AP) to Bregma) using a sterile 1 mm diameter micropuncher. Dissected AC tissue was then transferred to ice-cold centrifuge tubes and stored at −80 °C until processed for nuclei isolation.

### Nuclei isolation and fixation

Prior to performing nuclei fixation for the Parse Biosciences fixation protocol, nuclei from the AC were isolated by adapting the published 10X Genomics protocol for ‘Isolation of Nuclei for Single Cell RNA Sequencing’. Prior to and during nuclei isolation, all tools and buffers were kept chilled on ice. Frozen micropunches from two rats were combined and briefly thawed on wet ice. Then, the micropunches were transferred to chilled lysis buffer (10 mM Tris-HCl, 10 mM NaCl, 3 mM MgCl_2_, 0,1% Nonidet™ P40), dissociated until homogeneous with a pestle, and incubated on ice for 10 min. Micropunches were passed through a pre-chilled nuclei isolation column and centrifuged at 500xg for 3 min at 4°C to pellet nuclei. Nuclei-pellets were then washed in a ‘nuclei wash and resuspension buffer’ [1× PBS (phosphate-buffered saline), 1% BSA (bovine serum albumin), 0.2 U/μl RNase inhibitor], filtered, and pelleted again. Resuspended nuclei were filtered twice through a 30 μm strainer and counted before proceeding to the nuclei fixation protocol (Parse Biosciences V2). Nuclei were then partitioned to ensure there were less than 4 million nuclei in each aliquot and centrifuged at 200 x g for 10 min at 4°C. Pellets were resuspended in 750 µL nuclei buffer containing 0.75% BSA (Parse Biosciences) and then fixed using the Parse Biosciences Nuclei Fixation kit (Catalog # ECF2003, Parse Biosciences, Seattle, WA). Briefly, samples were passed through a 40 µm filter, incubated in nuclei fixation solution for 10 minutes on ice, followed by incubation in nuclei permeabilization solution. 4 mL of nuclei neutralization buffer was added to each sample, and they were centrifuged at 500xg for 10 minutes at 4°C. Pellets were resuspended in nuclei buffer with DMSO (1:20), frozen, and stored at −80°C until library preparation.

### Barcoding and Library Preparation

In advance of nuclei barcoding and library preparation, fixed nuclei were removed from −80°C, thawed in a water bath set to 37°C, placed on ice, and counted. Barcoded single-cell libraries were prepared from fixed single-cell suspensions using Evercode Whole Transcriptome v.2 (Catalog # ECW02030, Parse Biosciences, Seattle, WA) following the manufacturer’s instructions. Briefly, fixed nuclei were subjected to four rounds of barcoding in four reaction plates to generate single nuclei RNASeq libraries. Nucleus-specific barcodes are added on Round 1 of barcoding according to the manufacturer’s instructions (Parse Biosciences, Seattle, WA). In this round, PolyT primers and random hexamers bind RNA transcripts to generate barcoded cDNA in the presence of reverse transcriptase. At the end of the reaction, nuclei from all wells of the Round 1 plate were pooled and combined with a ligation mix. 40ul of nuclei suspension was then added to each well of the Round 2 barcoding plate and incubated. After stopping, the Round 2 ligation nuclei were again pooled and filtered through a 40 µm strainer. The filtered nuclei were subjected to a third round of barcoding by ligation in the Round 3 plate. After adding the Round 3 stop mix, nuclei were pooled again and filtered through a 40µm strainer, washed, and counted. The pooled nuclei were distributed into multiple sublibraries based on the manufacturer’s instructions. cDNA isolated from each sublibrary was subjected to generation of Illumina compatible sequencing library. The fourth barcode was incorporated during PCR amplification of the sequencing library.

### Single-nuclei RNA sequencing (Parse v2) and data processing

Cells were captured using the Parse Split-seq method (Parse Biosciences, Seattle, WA). Sublibraries were pooled at an equimolar ratio and sequenced on an Illumina NovaSeq 6000 platform (Illumina, Inc. San Diego, CA) with a sequencing configuration of 74-6-0-86 to generate ∼50k reads per cell. Raw reads were barcode deconvoluted and aligned to the reference genome (rnor7) via split-pipe (v1.04). All subsequent processing was performed using the Seurat package within R (v4.3.0)^85^.

### QC and clustering

Ambient RNA was reduced (SoupX v1.6.2), low-quality cells (cells with a percentage of reads of mitochondrial origin >10%, with a percentage of reads of ribosomal origin >45%, with <100 genes, with >7000 genes, <1000 total reads) were filtered from the dataset, and read counts were normalized using the scTransform method^86,87^.The sample was clustered via UMAP^88^ according to nearest neighbors. One cluster had canonical markers indicative of being derived from the adjacent hippocampus. The Allen Brain Atlas tissue profiles for the hippocampus^89–91^ were used to calculate signature scores per cluster to verify hippocampal cell contamination, and the cluster was subsequently removed from further analysis (**Supplemental Figure 1**).

Marker genes per cluster were identified using the FindMarkers function in Seurat with the Wilcoxon rank sum test and only positive markers flag to identify all genes significant per respective cluster versus all other cells. Gene Ontology analysis (GO)^92,93^ was calculated using an over-representation test within clusterProfiler (v4.6.2) using all significant markers (padj < 0.05) per respective cluster. Cell annotations were generated based on canonical and previously reported markers^5,6,15,36,40,42,94^ as compared to the gene expression profiles per cluster (**Supplemental Table 1)**.

## Data availability

The snRNA-Seq data generated in this study has been deposited to Gene Expression Omnibus (GEO) under the entry GSE262970 (https://www.ncbi.nlm.nih.gov/geo/query/acc.cgi?acc=GSE262970). Further inquiries can be directed to the corresponding author.

## Code Availability

The code for data analysis is available from the corresponding author on request.

## Acknowledgments

We thank our collaborators the Immune Monitoring Shared Resource team at Rutgers, Cancer Institute of New Jersey for assistance with the processing of the tissue sample; and the Genomics Center team at Rutgers New Jersey Medical School for assistance with library preparation and sequencing.

## Author contributions

Conceptualization: A.G.A., A.L. and K.M.B.; Methodology: A.G.A., K.M.B.; Investigation: A.G.A.; Visualization: A.L.; Funding acquisition: K.M.B.; Supervision: K.M.B.; Writing – original draft: A.G.A.; Writing—review and editing: A.G.A., A.L. and K.M.B. All authors read and approved the manuscript.

## Funding

This work was supported by the National Institutes of Health, National Institute of Deafness and Communication Disorders [R01-DC018561 to K.M.B.]

## Competing interests

The authors declare no competing interests.

## Supplementary Information

**Supplementary Table 1:** Gene lists for transcriptomic cell types.

**Supplementary Figure 1.**
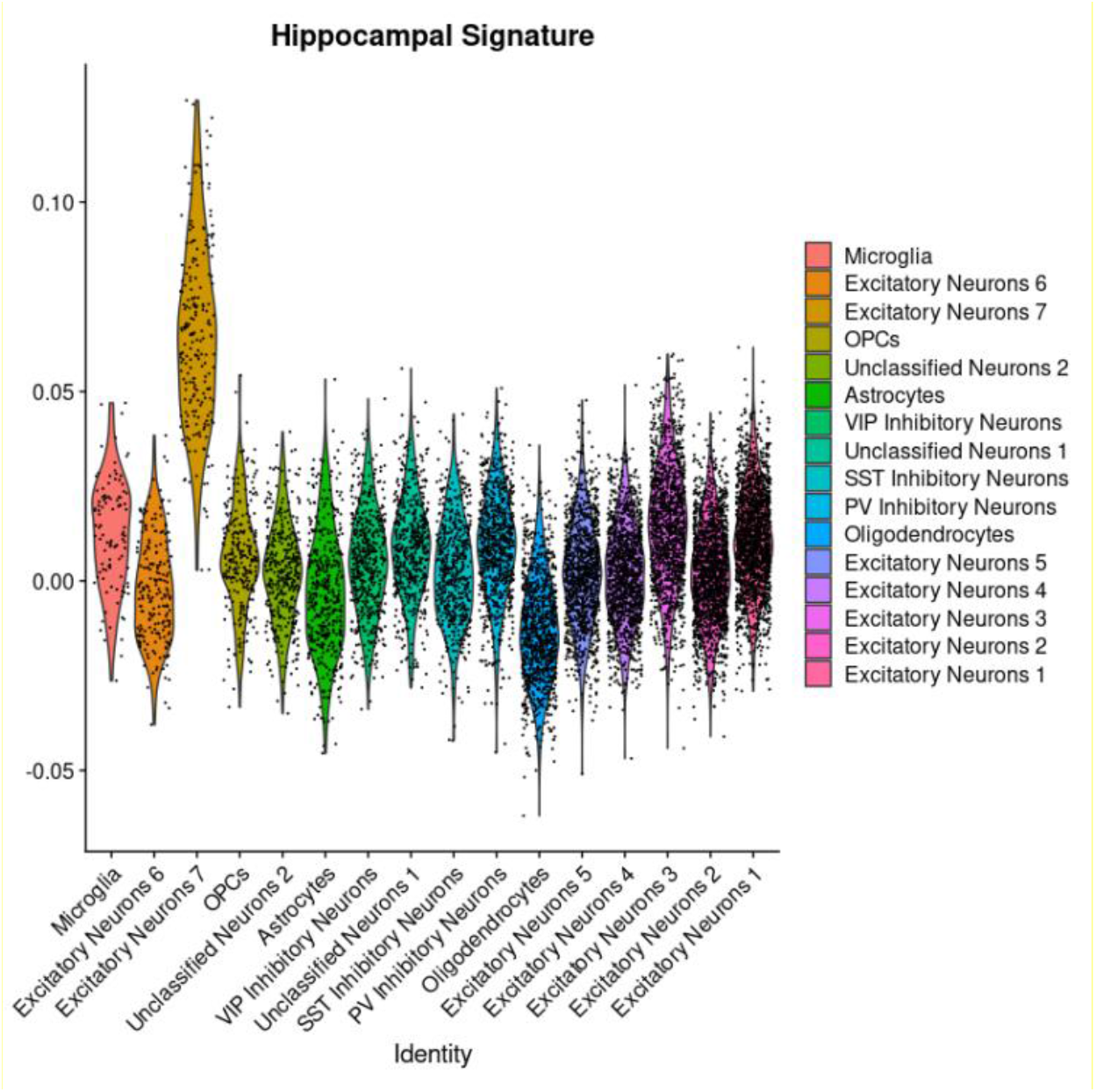
The hippocampal molecular signature across all identified clusters. Violin plots displaying scoring of the power relative to the other cell clusters using gene signatures for the hippocampus.

## Notes

https://www.ncbi.nlm.nih.gov/geo/query/acc.cgi?acc=GSE262970

### Competing Interest Statement

The authors have declared no competing interest.

